# Growth performance and nutritional composition of *Clarias gariepinus* in flood-and-drain aquaponics coupled with three species of leafy vegetable

**DOI:** 10.1101/2022.10.06.511190

**Authors:** Gbolaga O. Olanrewaju, David D. Sarpong, Abiola O. Aremu, Elizabeth O. Ade-Ademilua

## Abstract

Recirculating aquaponics integrate aquaculture and hydroponics with the aid of microorganisms to ensure a sustainable supply of fish and vegetables. In this study, we designed and constructed a flood-and-drain aquaponic system with *Clarias gariepinus* as the aquaculture and *Celosia argentea, Corchorus olitorius*, and *Ocimum gratissimum* as plant components. Nitrogenous waste from the aquaculture unit was circulated to the plant growth beds, which served as bio-oxidizers of toxic ammonia to nitrate and returned less toxic water back to the aquaculture unit. An evaluation of the growth parameters of *C. gariepinus* in aquaponics and control fish tanks revealed that aquaponics-raised *C. gariepinus* gained an additional 205.6% of their initial mean weight, whereas those grown in the control fish tank gained an additional 182.2% of their initial mean weight. The majority (37.5%) of aquaponics-raised *C. gariepinus* weighed 750 g -1 kg, whereas the highest percentage of *C. gariepinus* raised in the control fish tank (23.4%) weighed 500 g - 700 g. *C. gariepinus* raised in aquaponics had significantly higher gross feed conversion efficiency and protein efficiency ratio than those raised in the control tank. The mortality rate in the aquaponic fish tank was 0% compared to the 11.43% mortality rate in the control fish tank. There was no significant difference in the nutritional composition of *C. gariepinus* raised in either tank; however, the aquaponic fish tank had a higher nitrogen retention rate. This study showed that *C. gariepinus* raised in aquaponics had better biomass accumulation than those raised in conventional fishponds.

## Introduction

Global fish consumption, with a yearly growth of 3.1%, outpaces the human population expansion rate of 1.6% and 1.1% global meat consumption rates, with aquaculture accounting for 52% of the global consumption of fish (OECD-FAO, 2020). Global phenomena, such as population explosion and rise in Earth’s temperature, with their attendant consequences, remain a challenge to global sustainable fish production. Fish farming faces the challenge of improving net agricultural productivity to sustainably meet food demand by current and future projected populations (Ma et al., 2022), while simultaneously reducing the environmental footprint (Zhou et al., 2022). The harsh realities of urbanization, loss of agricultural land (Carballeira Braña et al., 2021) and agricultural water sources (Schönenberger et al., 2022) make the adoption of smart urban fish farming a viable option for sustainable fish production. Aquaponics, a hydroponic-aquaculture complex, provides an economical and smart alternative for fish production when land and water resources are limited. This environmentally friendly system (David et al., 2022) mimics natural nutrient cycling (nitrogen cycle) by recruiting aquatic animals (fish), plants, and microorganisms in a controlled manner for the sustainable simultaneous production of plants and fish. The microorganism (bacterial) component converts waste from aquaculture into nutrients that are readily available for uptake by the plants. The process of waste conversion is known as nitrification, and the cycle of this process is known as the nitrogen cycle. In this cycle, nitrogen exists in three main forms: ammonia (NH_4_^+^ or NH_3_), nitrite (NO_2_^-^), and nitrate (NO_3_^-^).

An explanation for the nitrogen cycle in fish tanks was proposed by Bik et al. (2019), who pointed out that ammonia, which is the main excretion product from fish, can be toxic to fish at very low levels. During nitrification, certain autotrophic bacteria (primarily *Nitrosomona*) oxidize ammonia to nitrite, whereas Nitrobacter oxidizes nitrite to nitrate. These bacteria colonize the surfaces of aquatic systems. Nitrifying bacteria thrive optimally at temperatures between 77 and 86 °F (Moloantoa et al., 2022). When temperatures fall below 32 °F or rise above 120 °F, these bacteria begin to die off, and aquatic system nitrification is halted. Another factor that must be accounted for in aquaculture is the water pH. Nitrifying bacteria thrive at pH between 7.3 and 7.5(Taylor et al., 2017). When the pH fell below 6.0, nitrification ceased and the system became toxic. The dissolved oxygen level, total dissolved solutes, and other water parameters must be examined and found to be conducive to the survival of these bacterial communities.

*Clarias gariepinus* (African sharp-tooth catfish) is a catfish species that belongs to the family Clariidae (air-breathing catfish). They are found throughout Africa and the Middle East and live in freshwater lakes, rivers, and swamps, as well as in human-made habitats, such as oxidation ponds or even urban sewage systems (Schoch et al., 2020). African catfish was introduced worldwide in the early 1980s for aquaculture. Therefore, it is found in countries far outside its natural habitat, such as Brazil, Vietnam, Indonesia, and India. African catfish is a large, eel-like fish, usually with a dark gray or black coloration on the back, fading to a white belly. *C. gariepinus* has an average adult length of 1-1.5 m and, reaches a maximum length of 1.7 m and weighs up to 60 kg. It is a nocturnal fish, like many catfish, and feeds on living as well as dead animal matter. Owing to its wide mouth, it can wholly swallow relatively large prey. It is known to consume large water birds such as the common moorhen (Afia & David, 2018). The choice of *Clarias gariepinus* for this study was based on its prevalent global consumption as food (Dauda et al., 2018), uncomplicated breeding, and excellent feed conversion ratio (Knaus et al., 2020).

Several studies have proposed and designed various forms of aquaponics (Kotzen et al., 2019), involving different species of aquaculture and plants. However, based on careful consideration of the pros and cons of different aquaponic system designs and the limited availability of financial and material resources, this research utilized a media-based design, also known as the flood-and-drain aquaponic system (Figure 1). The African catfish, *Clarias gariepinus*, was the preferred fish for the aquaculture unit, while three common vegetables in Nigeria, *Celosia argentea, Corchorus olitorius, and Ocimum gratissimum*, were the preferred plants for the hydroponic unit based on their food and medicinal values (Guzzetti et al., 2021; Nganteng et al., 2022). This study aimed to compare the growth parameters of *C. gariepinus* raised in constructed recirculatory aquaponics to those of *C. gariepinus* raised in conventional fishponds.

**Figure 1:**
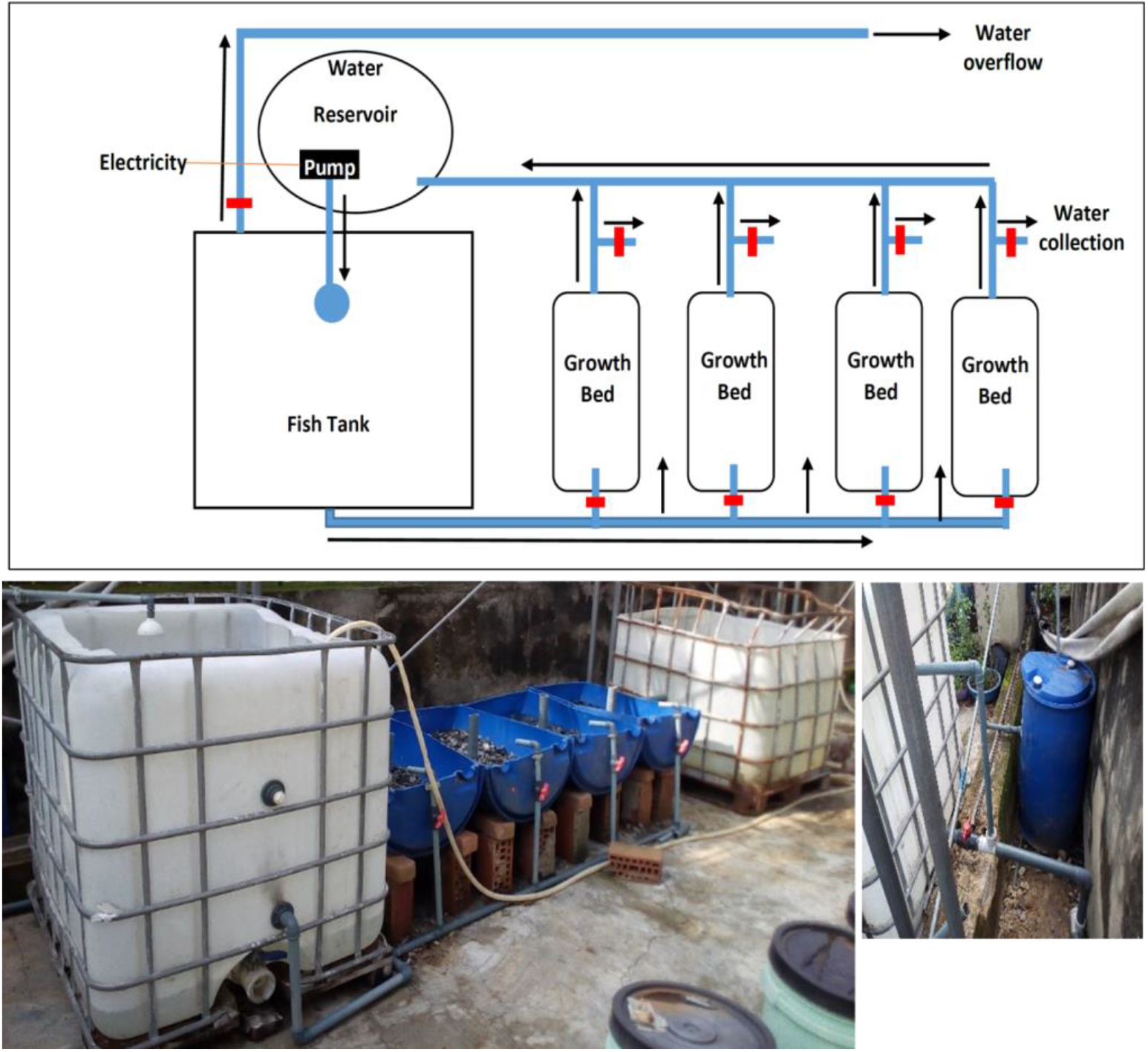
The aquaponics design. The aquaponics block design (top) and constructed aquaponics (bottom). The aquaponics system has 3 compartments which are the fish tank that contained *Clarias gariepinus*, the growth beds which contained four compartments filled with granite substrate for plant growth, and the water reservoir which receives nitrified water from the growth bed and returns it to the fish tank. Each of the compartments contains a water valve (in red) which controls the flow of water in and out of the compartment. The compartments were placed on different topography to enable water movement under the influence of gravity and reduce power consumption by the installed water pump. Water returned to the fish tank was manually oxygenated by using a sprinkler head dispenser.

## MATERIALS AND METHOD

### Design and construction of the aquaponics

The designed and constructed flood-and-drain aquaponic system was modeled after the FAO (Somerville et al., 2014) design template, with some modifications to reduce system power consumption using gravity-assisted water movement (Figure 1). The system consists of three units: an aquaponic fish tank, plant growth bed, and recycled water reservoir. The plumbing connection of the three units was established at varying heights to allow unidirectional water movement from the fish tank back to the fish tank via plant growth beds and recycled water reservoir. *Celosia argentea, Corchorus olitorius*, and *Ocimum gratissimum* were cultivated in the growth bed as plant components of the aquaponics. The aquaponic fish tank had an overflow release valve, and water was returned to it with a sprinkler head to allow oxygen to mix with the returned water (Figure 5). The control fish tank, a conventional pond, was set at the same location, and both were supplied with water from the University of Lagos.

### Stocking and feeding regime of *Clarias gariepinus*

One hundred and forty (140) pieces of 10 weeks old juvenile African sharp-tooth catfish, *Clarias gariepinus*, were purchased from the Aquaculture Unit, Department of Marine Science, University of Lagos. At stocking time, the juvenile catfish had a mean weight and height of 14.3 ± 1.2 g and 5.8 ± 0.8 cm respectively and were sorted into aquaponics and control fish tanks 2 weeks after stocking. This was done to cushion the effect of transportation stress on fish. At the commencement of the Aquaponic cycling, the mean weight of fishes in the aquaponic and conventional ponds were 87.6 + 5.3 g and 86.8 + 4.6 g respectively, while the mean height was 14.2 + 1.4 cm and 13.9 + 0.6 cm respectively. Growth measurement started an additional 2 weeks after the initialization of the aquaponics. Fish in both aquaponics and control fish tanks were fed copper feed (with the following nutritional composition: crude protein (40%), crude fat (9%), crude fiber (3.5%), ash (7%), total phosphorus (1%), lysine (2%), methionine and cysteine (1.16%), vitamins A, D2, E, and C, and antioxidants. The fish were fed twice daily, and the feed pellet size was increased at intervals of 3 weeks.

### Fish growth and nutrient utilization parameters

#### Mean weight

The initial weight of the fish in each tank was determined at the beginning of the experiment and recorded as a reference. The fish were weighed weekly using a weighing scale (Camry EK5055, Max. 5kg/11lb d=1g/0.05oz).

#### Mean Weight Gain (MWG)

The MWG was calculated using the formula below

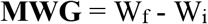

where Wf is the final average weight and Wi is the initial average weight (g).

#### Mean Length Gain (MLG)

The MLG was calculated as:

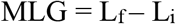

where L_f_ is the final average length (cm), and L_i_ is the initial average length (cm).

#### Weight-Length Relationship (WLR)

This was expressed as ***W****= aL*^*b*^ (Song et al., 2022)

where W= weight in grams, L is the length of the fish (cm), a is a constant (intercept), and b is a constant (slope of regression).

The LWR equation was transformed to its natural log prior to its correlation analysis.

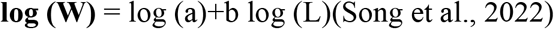

#### Feed Conversion Ratio (FCR)

This is the amount of feed per unit weight that the fish were able to convert into flesh.

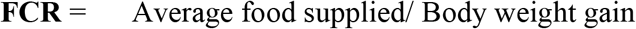

#### Protein Intake (PI)

Protein intake was calculated using the formula below

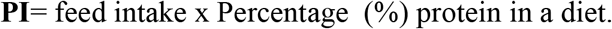

#### Protein Efficiency Ratio (PER)

This was calculated from the relationship between the increment in the weight of fish (i.e., weight gain of fish) and the amount of protein consumed.

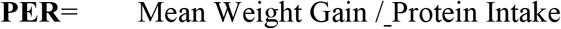

#### Gross Food Conversion Efficiency (GFCE)

This was calculated as the reciprocal of the FCR expressed as a percentage.

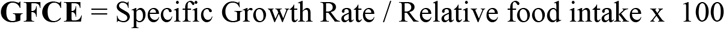

### Proximate analysis

The proximate composition of fish samples from the aquaponic and control fish tanks was determined according to the AOAC(Association of Official Analytical, 2005) protocol at the Department of Animal Sciences of the University of Ibadan, Oyo State, Nigeria. The moisture content was determined by drying the samples overnight at 105 °C for 2 h and 30 min. The crude protein content was determined using the Kjeldahi method. The fat content was determined using the Soxhlet method described by (López-Bascón & Luque de Castro, 2020) and the ash content was determined by ashing at 500 °C until fully ashed. Carbohydrate content was determined by calculating the difference; that is, the sum total of moisture, protein, fat, and ash content was subtracted from 100.

### Determination of water pH, TDS and nitrogen dynamics

The total dissolved solutes (TDS) and pH of the water were measured weekly using a handheld digital API Master TDS and pH monitor. The concentration of non-ionized ammonia in the collected water samples was determined by the salicylate method(Giner-Sanz et al., 2022), nitrite concentration by the diazotization method (el hani et al., 2022) and nitrate concentration by the Szechrome NAS reagent (Taylor et al., 2010).

## RESULTS

Several studies have reported that the physicochemical conditions of the aquatic environment have profound effects on fish growth physiology (Stavrescu-Bedivan et al., 2016; Wang et al., 2022), nutrient conversion efficiency, and the capacity to adapt to the environment (Calduch-Giner et al., 2022). To understand how our designed aquaponic system environment affected the growth and development of *Clarias gariepinus* (African catfish), we measured *C. gariepinus* physiological growth parameters over 10 weeks.

### Analysis of *Clarias gariepinus* weight and length in aquaponics

Despite starting both aquacultural systems with fishes having similar weights (aquaponics, 87.6±0.4g and control fish tank, 86.8±0.3), *C. gariepinus* raised in the aquaponics had significantly higher weights (p<0.05) than those raised in the control fish tank (Figure 2a) through the 4th week to 10th week of observation. In comparison to the control fish tank, with 4.8% of harvested fish weighing more than 1 kg, 20.3% of *C. gariepinus* harvested from aquaponics weighed more than 1 kg (Table 1). The highest percentage (37.5%) of aquaponics-raised *C. gariepinus* weighed between 750 g -1 kg, while the highest percentage of *C. gariepinus* raised in the control fish tank weighed 500 g -700 g. Only 18.8% of *C. gariepinus* raised in aquaponics weighed below 500 g, whereas 29% of *C. gariepinus* raised in the control fish tank weighed below 500 g (Table 1).

**Table 1:** Weight classification of *Clarias gariepinus* from the aquaponics and control fish tank at full marketable age maturity

| Weight | % Number of fishes |  |
| --- | --- | --- |
|  | Aquaponics | Conventional culture tank |
| Extra big (> 1kg) | 20.3 | 4.8 |
| Big (750g - 1kg) | 37.5 | 29 |
| Medium (500g–700g) | 23.4 | 37.1 |
| Small (< 500g) | 18.8 | 29 |

**Figure 2:**
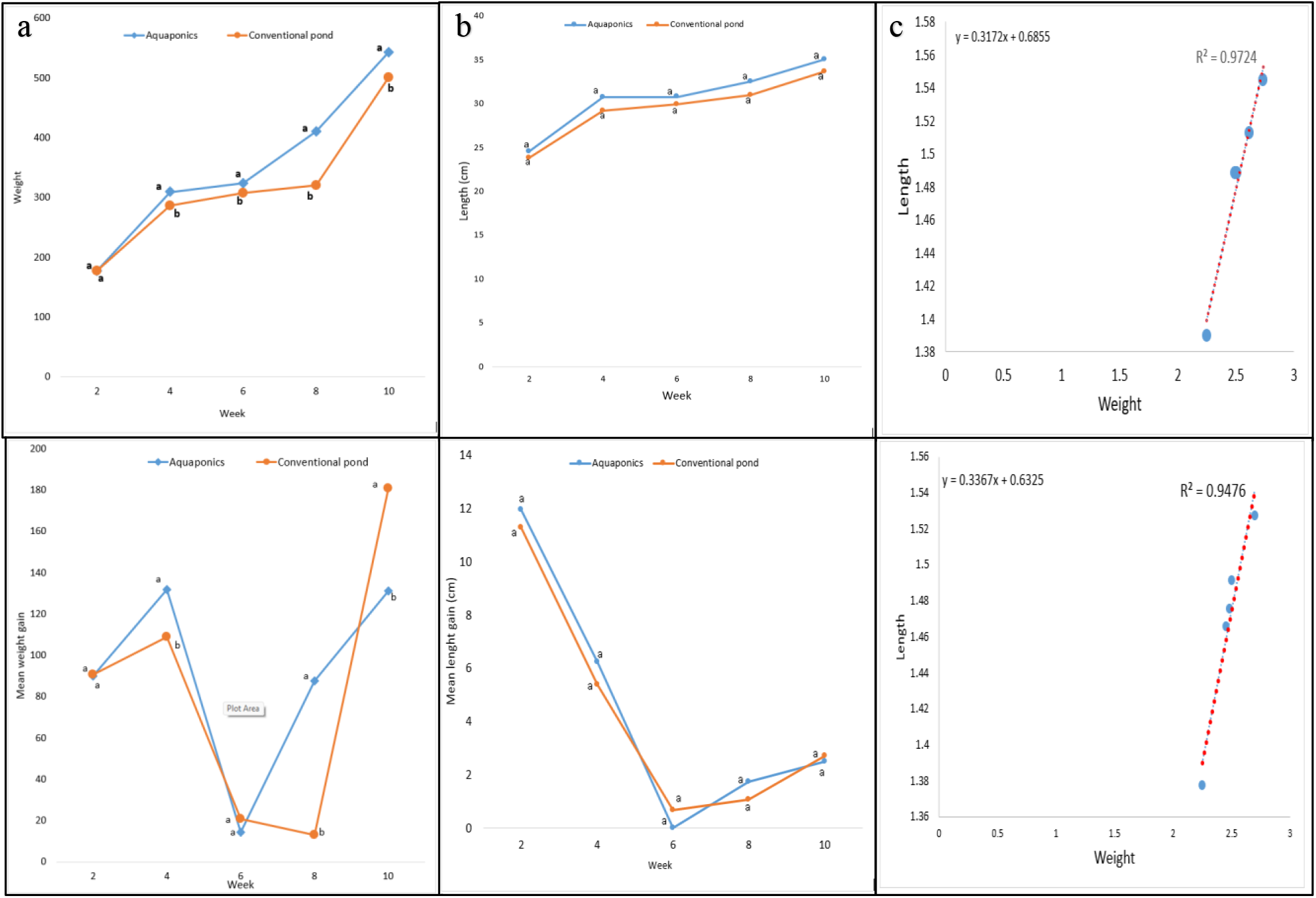
Weight and Length parameters of *C. gariepinus*. Plotted means, in same week and on the same horizontal axis, represented with different letters are significantly different at p= 0.05 **a**. The mean weight (top) and the mean weight gained per 2 weeks (bottom) of *C. gariepinus* in aquaponics and control fish tanks, across the weeks of observation. **b**. The mean length (top) and mean length gain of *C. gariepinus* in aquaponics and control fish tanks, across the weeks of observation **c**. Correlation plot of the weight-length relationship of *C. gariepinus* in aquaponics (top) and control fish tanks (bottom). In **a-b**, blue lines indicate the aquaponics while orange lines indicate the control fish tank, referred to as the conventional pond.

The weight gain by *C. gariepinus* raised in aquaponics was significantly higher (p<0.05) than those in the control fish tank at weeks 2 and 8. However, *C. gariepinus* raised in the control fish tank gained significantly more weight between weeks 8 and 10 than those grown in the aquaponics (Figure 2a). At the end of the study, *C. gariepinus* raised in aquaponics gained an additional 205.6% of their initial mean weight, whereas those grown in the control fish tank gained an additional 182.2% of their initial mean weight. Throughout the study, *C. gariepinus* raised in aquaponics were consistently longer than those in the control fish tank. There was a decrease in the length gained in two weeks in both fish tanks, with aquaponics-raised *C. gariepinus* gaining more length in two weeks than *C. gariepinus* in the control fish tank. However, the differences in length and mean length gained per 14 days were not significant (Figure 2b). *C. gariepinus* raised in aquaponics had a weight-length relationship of W =0.32L^0.69^, whereas those raised in the control fish tank had a weight-length relationship expressed as W= 0.34L^0.63^. There was a strong positive correlation between weight and length in the aquaponic (R^2^ =0.97) and the control fish tank (R^2^= 0.95) (Figure 2c).

### Feed conversion ratio (FCR), gross feed conversion efficiency (GFCE) and protein efficiency ratio (PER)

The feed conversion ratio, expressed as the ratio of average feed supplied to weight gained by the fish at 14 days (2 weeks), was not significantly different (p<0.05) between *C. gariepinus* raised in aquaponics and those raised in the control tank in the 2nd through 4th week and at the 10th week of the study. However, there was a significant spike in the feed conversion ratio of *C. gariepinus* raised in aquaponics in the 6th week while those raised in the control tank had a similar significant spike in the 8th week. Implicated by the FCR, *C. gariepinus* raised in aquaponics had above 60% and 30% gross feed conversion efficiency at weeks 4 and 8, respectively, which were significantly higher than those observed in the control fish tank. However, *C. gariepinus* raised in the control fish tank had a significantly higher feed conversion efficiency in the 10th week compared to those raised in aquaponics (Figure 3a). Cumulatively, *C. gariepinus* raised in aquaponics were more efficient in converting feed to biomass. The distribution of the protein efficiency ratio (PER), calculated as the ratio of the mean weight gained by fish to protein intake in 14 days, was similar to that of the gross feed conversion efficiency. A significant difference in PER was observed at weeks 4, 8, and 10, with *C. gariepinus* raised in aquaponics having a higher PER than those raised in the control tank in weeks 4 and 8, the PER was vice-versa in the 10th week (Figure 3b).

**Figure 3.**
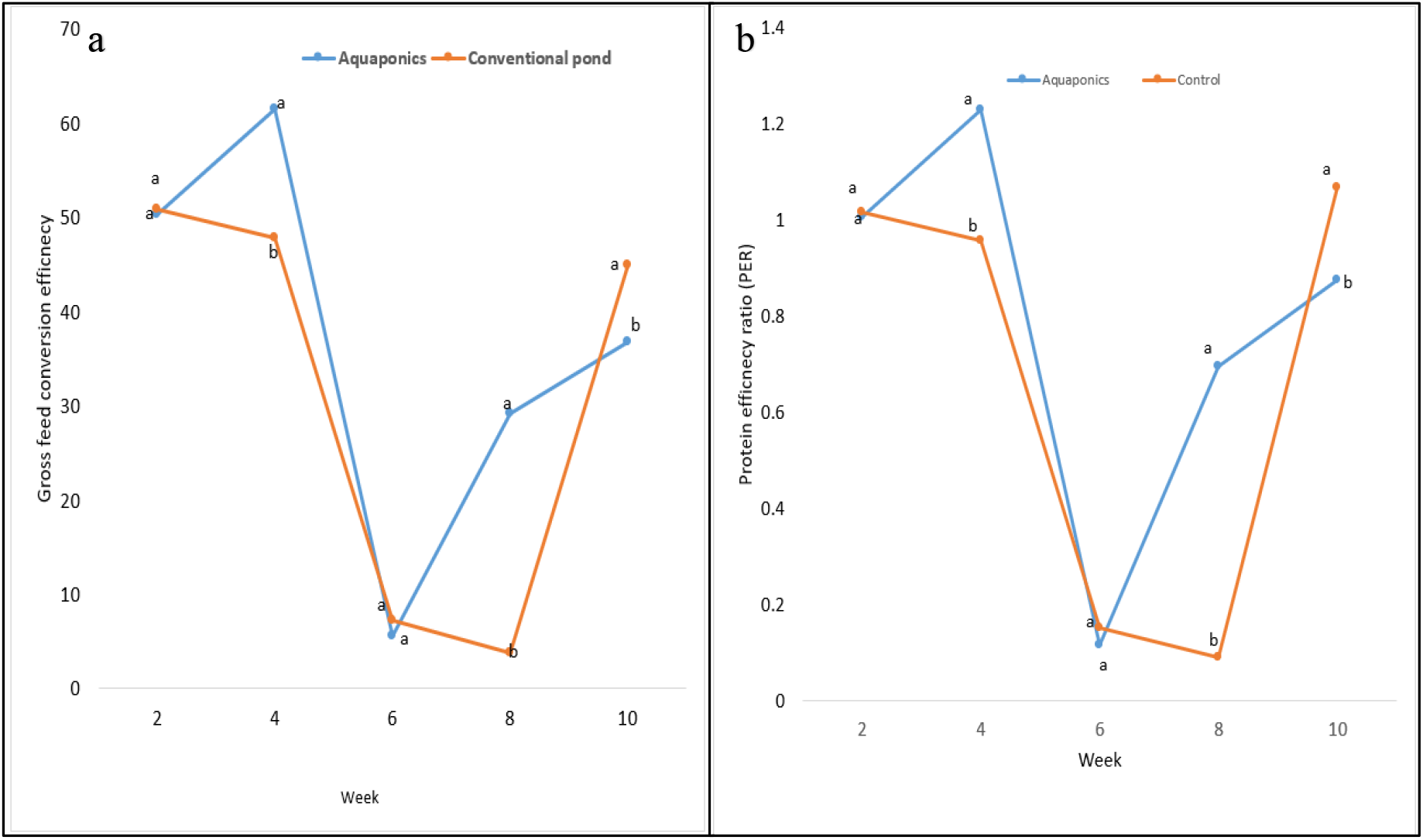
Gross feed conversion efficiency (GFCE) and Protein Efficiency Ratio (PER). **a**. The GFCE of *C. gariepinus* raised in aquaponics and control fish tank (conventional fishpond) over the 10 weeks of observation. Plotted means, in the same week and on the same horizontal axis, represented with different letters are significantly different at p= 0.05. **b**. The Protein efficiency ratio (PER) of *C. gariepinus* in aquaponics and control fish tank (conventional fishpond) over the 10 weeks of observation. Plotted means, in the same week and on the same horizontal axis, represented with different letters are significantly different at p= 0.05. In **a-b**, the blue line indicates aquaponics while the orange colored line indicates the control fish tank (conventional pond).

### Mortality rate of *Clarias gariepinus*

*C. gariepinus* raised in aquaponics showed no mortality throughout the study period, in contrast to the frequency of mortality observed in *C. gariepinus* raised in the control fish tank, which lost 11.43% of its total population by the end of week 10 (Figure 4a).

**Figure 4.**
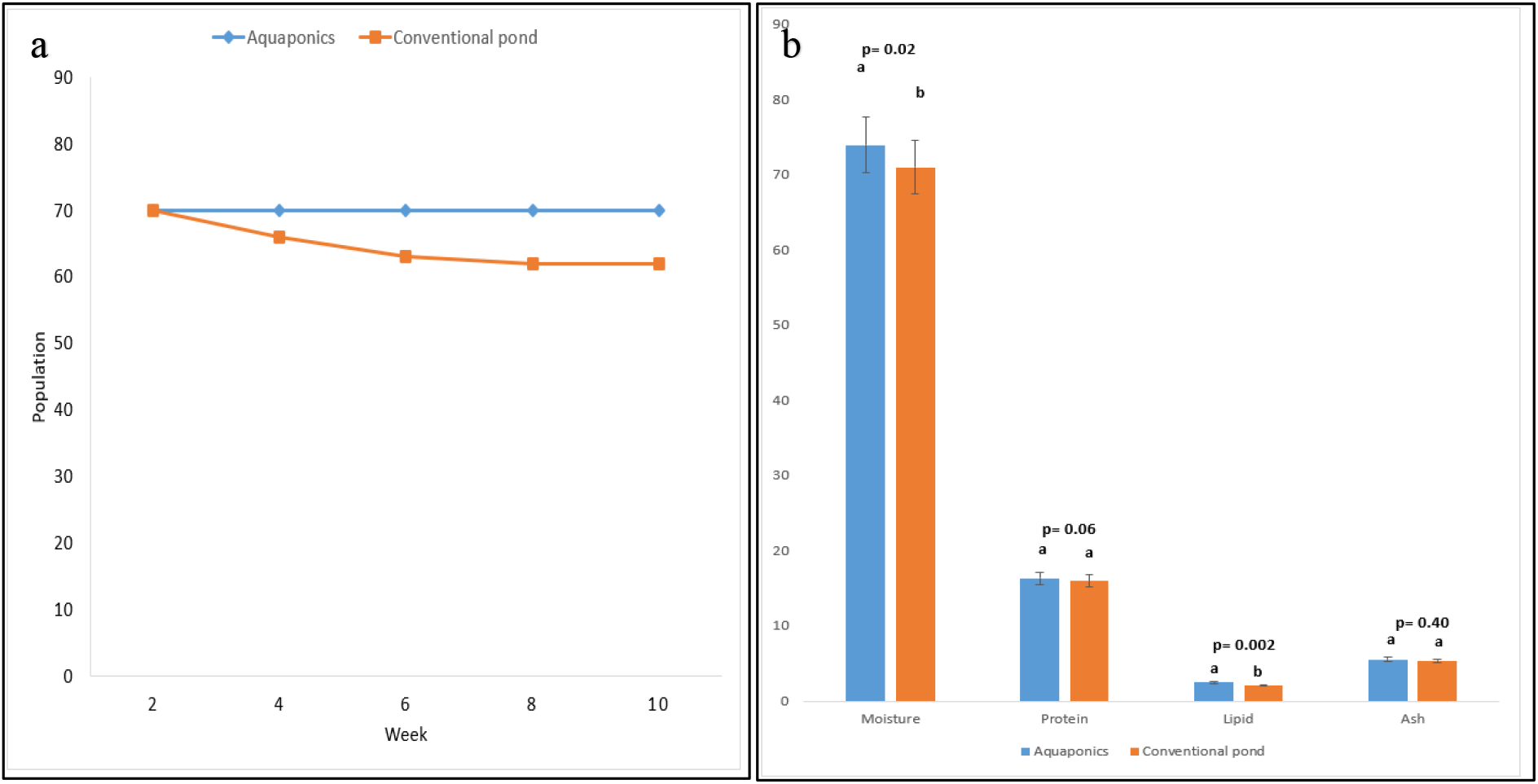
Mortality and Nutritional composition of *C*. gariepinus **a**. The mortality of *C. gariepinus* raised in aquaponics and control fish tank (conventional fishpond) over the 10 weeks of observation. Plotted means, in the same week and on the same horizontal axis, represented with different letters are significantly different at p= 0.05. **b**. The nutritional composition of *C. gariepinus* in aquaponics and control fish tank (conventional fishpond), as determined by proximate analysis, over the 10 weeks of observation. Column bars representing the same nutrient and on the same horizontal axis, represented with different letters are significantly different at p= 0.05. In **a-b**, the blue line indicates aquaponics while the orange colored line indicates the control fish tank (conventional pond).

### Nutritional composition

Proximate analysis of the nutritional composition of *C. gariepinus* raised in aquaponics and those raised in the control fish tank revealed that *C. gariepinus* raised in aquaponics had significantly higher moisture and lipid contents than those raised in the control tank. The %protein and %ash contents were not significantly different between the fish from both tanks (Figure 4b).

### Nitrogen dynamics, Total Dissolved Solids and pH in fish tanks

Except for the 2nd week when aquaponics raised *C. gariepinus* produced significantly lower (p<0.05) concentrations of ammonia (mg/l) than those raised in the control fish tank, the production of ammonia was not significantly different between *C. gariepinus* raised in the aquaponics and those raised in the control fish tank throughout the study period (Table 2). However, nitrate concentrations (mg/l) were significantly higher (p<0.05) in the aquaponics tank than in the control fish tank in the 2nd through the 8th week, while no significant difference was observed in nitrate concentration between the two ponds in the 10th week (Table 2). The total dissolved solids (TDS) were consistently higher in the aquaponic tank (Table 2) than in the control fish tank, which also had consistently higher TDS than the aquaponic recycled water reservoir. The pH in both ponds was maintained at a range of 6.5 -7.8 (Table 3).

**Table 2:**
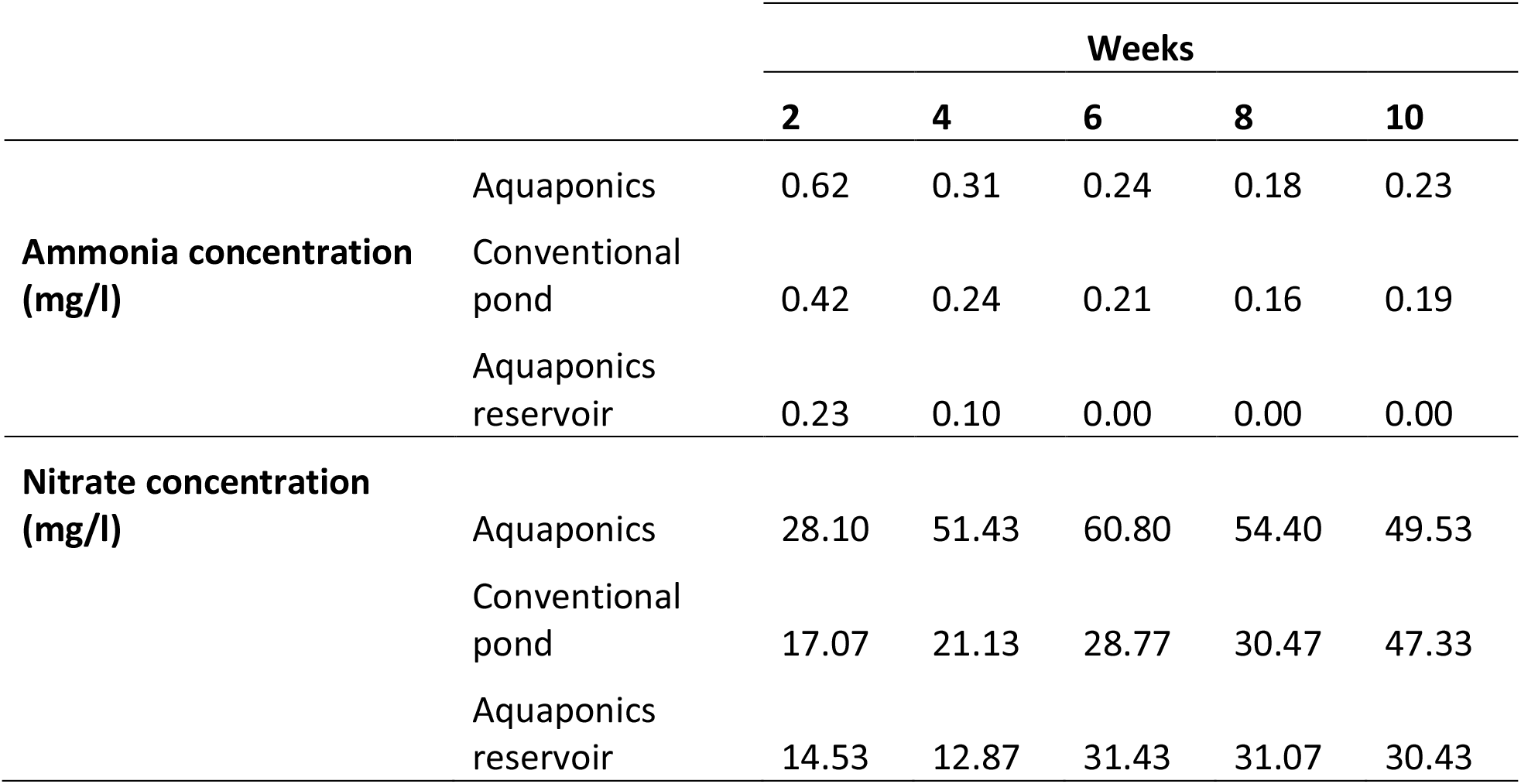
Ammonia and Nitrite concentration (mg/l) in the aquaponics, control fish tank and the aquaponics reservoir

**Table 3:** System pH and Total dissolved solids (mg/l) in the aquaponics, control fish tank and the aquaponics reservoir

|  |  | Weeks |  |  |  |  |  |  |  |  |  |
| --- | --- | --- | --- | --- | --- | --- | --- | --- | --- | --- | --- |
|  |  | 1 | 2 | 3 | 4 | 5 | 6 | 7 | 8 | 9 | 10 |
| **System pH** | Aquaponics | 7.8 | 7.6 | 6.9 | 6.9 | 6.8 | 6.8 | 7.3 | 6.9 | 6.8 | 6.3 |
|  | Conventional pond | 7.4 | 7.6 | 7.1 | 6.9 | 6.8 | 6.9 | 7.4 | 7.8 | 7.4 | 7.3 |
|  | Aquaponics reservoir | 7.4 | 6.9 | 7.1 | 6.8 | 6.9 | 6.8 | 6.9 | 6.5 | 6.4 | 6.3 |
| **System TDS (mg/l)** | Aquaponics | 549 | 654 | 679 | 736 | 846 | 852 | 538 | 683 | 645 | 682 |
|  | Conventional pond | 512 | 640 | 608 | 684 | 798 | 803 | 489 | 614 | 611 | 626 |
|  | Aquaponics reservoir | 485 | 598 | 520 | 648 | 704 | 783 | 448 | 601 | 586 | 541 |

## DISCUSSION

### *Clarias gariepinus* gained more biomass in the aquaponics

The basic idea behind the operation of aquaponics is the simultaneous cultivation of aquaculture and plants, where both are mutually dependent on each other for survival (Paudel, 2020), with the relationship maintained by the efficient establishment of the nitrogen cycle. Our results indicate that the aquaponic environment encourages the accumulation of biomass, which is reflected in the higher weights of *C. gariepinus* raised in the system compared to those raised in the conventional pond. *C. gariepinus* raised in the conventional pond appeared to have delayed the achievement of growth milestones compared to those in aquaponics (Figure 5). Aquaponics mimics a natural pond that often has its water diluted by flowing, rather than being completely replaced, as seen in conventional pond management. Changing pond water creates a significant change in environmental parameters for fish, which may create sudden biological shocks and difficulty in adaptation (Rytwinski et al., 2020). The downward steepness observed in the mean weight gain in both fish tanks between weeks 4 and 6 was a result of inappropriate feed pellet size fed to fish in both tanks. The feed pellet sizes introduced to the fish between those weeks of growth were larger than expected for the fish age (4 mm instead of 3 mm), which resulted in a deficiency in feed uptake by the fish. However, the aquaponics-raised *C. gariepinus* easily adjusted to their normal metabolism, as observed in their normalized weight gain trend, when the right feed size was introduced, compared with *C. gariepinus* raised in the control fish tank, which took more time to return to normal metabolism. This suggests that the aquaponic system increased the adaptive capability of *C. gariepinus*, and the fish could easily regain their normal metabolism once external stressors were removed. The weight-length relationship of fish in both tanks indicated allometric growth in weight (b <3). This suggests that the fish in both tanks did not have a proportionate increase in length with weight in a cubic form. However, the high correlation between weight and length in both fish tanks indicated a highly positive relationship between these parameters across the fish tanks. These results corroborate several other studies(Atique et al., 2022; Obirikorang et al., 2021; Oladimeji et al., 2020) that reported improved weight of fish raised in aquaponics compared to those raised in conventional fishponds.

**Figure 5.**
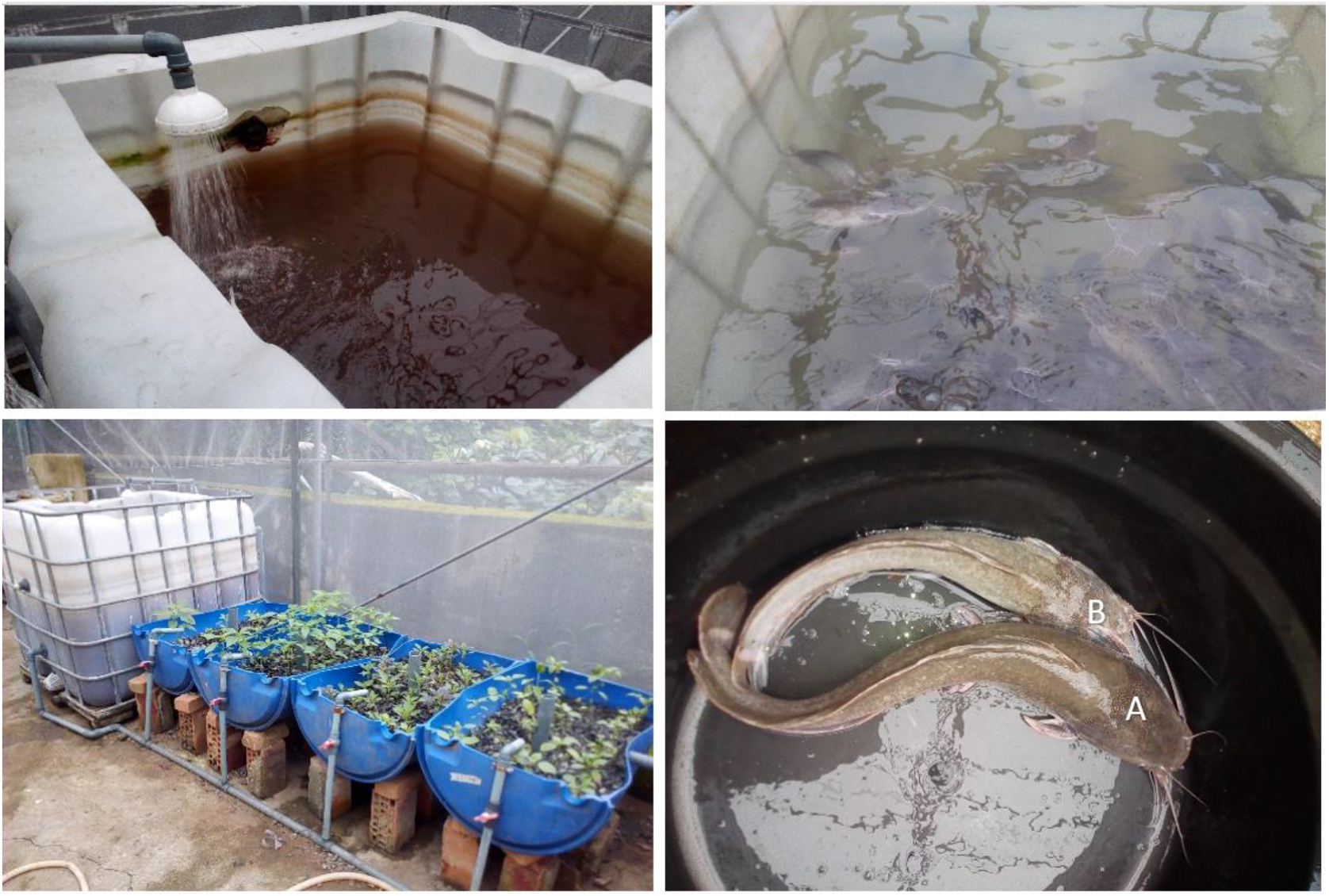
Overview of the aquaponics system operations and performance. **Left-top** is the brackish aquaponics pond with the sprinkler head releasing water into the aquaponics fish tank from the aquaponics reservoir. **Left-bottom** is a broad figure showing the plant growth bed and the aquaponics fish tank in operation. **Right-top** is the control fish tank after the water was newly changed. **Right-bottom** is a comparison of *C. gariepinus* from the aquaponics and the control fish tank at the 8th week. The letter A indicates aquaponics while B indicates the control fish tank.

### Aquaponics-raised *Clarias gariepinus* are more efficient at feed conversion

Environmental conditions have been reported to affect the feeding patterns, feed conversion efficiency, and feeding success of fish (Lopez-Betancur et al., 2020). The aquaponics environment improved the feeding efficiency of *C. gariepinus*, as observed in FCR, GFCE, and PER. Feeding dynamics in a fish tank start from the uptake of feeds by the fish, which Lopez-Betancur et al. (2020) reported to be lower in dark fish tanks than in light fish tanks, and he alluded this to increased visibility in the light fish tanks. However, our results do not agree with it, as water in the aquaponic fish tank appears brackish compared with the regularly changed water in the control fish tank. This study cannot convincingly ascertain whether feed uptake was higher in aquaponics, but this is likely to be the case. Studies have reported better feed conversion efficiency in Nile tilapia raised in aquaponics than in those raised in conventional fishponds (Amin et al., 2021; Shaw et al., 2022; Strand et al., 2007). An important observation in the aquaponic fish tank was the presence of slimy cyanobacteria (Figure 5), which increased with time and has been reported to significantly contribute to more than 10% of fish protein intake and affect their gut microbiome(Rosenau et al., 2021). Changes in the gut microbiome composition have profound effects on the physiological and molecular functions of *C. gariepinus* (Adejonwo et al., 2020). This study, therefore, suggests that *C. gariepinus* raised in aquaponics has access to sources of proteins other than administered feed, which is an advantage over the conventional pond system. This also affected the accuracy of PER estimation, as we could not effectively account for extra-feed protein intake. However, PER calculation based on weight gain and administered feed quantity indicated a better GFCE and PER in *C. gariepinus* raised in aquaponics compared to those raised in the conventional fishpond. A comparison of the nutritional content of *C. gariepinus* raised in both ponds however did not indicate a nutritional benefit of one over the other.

### Mortality and Nitrogenous dynamics in *Clarias gariepinus aquaponic*

The results of the analysis of *C. gariepinus* mortality in both systems further suggest that aquaponics improved fish adaptive capability, as zero mortality was observed in aquaponics compared to the 11.43% mortality observed in the control fish tank. Mortality observed in the control fish tank may be due to shocks introduced by the regular complete water-changing regime, which might lead to disruption of osmoregulation and functional metabolism. (Oladimeji et al. (2020) reported a significant improvement in the survival rate of *C. gariepinus* raised in pumpkin aquaponics compared to that of the control. The high mortality rate of *C. gariepinus* is a major source of revenue for farmers(Fernández Sánchez et al., 2022). The higher level of total dissolved solids in aquaponics was not surprising, as it is an active circulatory system with a flow-like water regime pattern, rather than a complete replacement of water, as in the conventional tank. However, this had no profound effect on the pH of the system, as the pH values in both the systems were similar. Nitrogen retention in the aquaponics was higher than that in the control tank, with an improved onset of nitrification. This is because of the presence of plants in the aquaponic system(Colt & Semmens, 2022).

## CONCLUSION

Aquaponics is an attractive agricultural concept with huge prospects for promoting sustainable agriculture compared to available conventional agricultural systems. This study provides insight into the growth performance of *C. gariepinus* in constructed low-power flood-and-drain aquaponics. The results of this study indicated an overall better yield of *C. gariepinus* raised in aquaponics, characterized by increased weight gain, increased feed and protein conversion efficiency, and reduced mortality.

## Supporting information

Raw data

## Authors’ contributions

The authors confirm the contribution to the paper as follows:

Study conception and design: GOO & EOA

Data collection: GOO & AOA

Analysis & interpretation of results: GOO, EOA, DDS &AOA

Draft manuscript preparation: GOO & DDS

All authors reviewed and approved the final version of the manuscript submitted

## Funding Statement

This research was jointly funded by the authors, and no external funding was received to conduct the research.

## Data access statement

All relevant data are within the paper and its supporting information files

## Competing interests

The authors declare that the research was conducted in the absence of any commercial or financial relationships that could be construed as potential conflicts of interest.

## Ethics approval

This study was approved by the ethics committee of the University of Lagos, Nigeria.

## Notes

### Competing Interest Statement

The authors have declared no competing interest.

